# Reference-guided automatic assembly of genomic tandem repeats with only HiFi and Hi-C data enables population-level analysis

**DOI:** 10.1101/2023.12.07.570710

**Authors:** Huaming Wen, Weihua Pan

## Abstract

The existing de novo methods of complete genome assembly are not able to generate large-scale pangenomes with complete assemblies due to the shortcomings such as requiring multiple types of sequencing data of high price, requiring large amount of manual curation, and not being able to achieve haplotype-resolved complete assembly of long tandem repeats in most situations. To solve this problem, in this study, we propose a new genome assembly mode called reference-guided assembly which relies on the reference information to recall the reads for complex genomic regions of interest and assembles them in *de novo*-like way. As a proof-of-concept, we developed an algorithm TRFill which can reassemble or fill the gaps of tandem repeats in chromosome-level assembly in either haploid or diploid way using only HiFi and Hi-C data. The experimental results on human centromeres and tomato subtelomeres show that TRFill successfully improved the completeness and correctness of about two thirds of the tested tandem repeat sequences. Furthermore, TRFill improved the completeness of subtelomeric tandem repeats by 50% in a recently published tomato pangenome, enabling a population-level analysis of the subtelomeric tandem repeats, which found the ‘local law of sequence similarity of tandem repeats’ providing theoretical basis for reference-guided assembly in turn.

## Introduction

For a long time, the reference sequences of main eukaryotic genomes are not complete, due to the missing of complex genomic regions such as tandem and interspersed repeats. The incomplete reference genomes impede functional, structural and evolutional studies of repetitive sequences^1, 2^ and may cause mistakes in reference-based data analysis such as data contamination^3^ and false-positive variant calls^4^. In recent years, with the advances in long-read sequencing technologies, especially the development of PacBio HiFi and ONT UL, the completeness of the eukaryotic reference genomes has been improved significantly. A number of corresponding assembly algorithms and tools have been developed^5–8^ and the T2T (Telomere-to-telomere) or near-T2T level reference genomes of at least dozens of species have been built^9–12^.

Although the technology has been available for generating a complete single genome, there are a couple of practical factors stopping building a population of complete genomes, leading to a very limited number of populational studies of repetitive sequences^13^. First, since the automatic assemblers (e.g., hifiasm^7^, hiflye^5^, hicannu^6^, verkko^8^) can only generate complete chromosomes in very few genomes, for most species, the complete genome cannot be built without enough manual process of filling the gaps in repetitive genomic regions which needs high professional techniques and is very time-consuming. Except for extremely important species like human, we believe it is almost impossible to gather a large number of experts in genome assembly and spend a large amount of time to finish the manual work for a population of dozens or even hundreds of genomes in any other species. Second, although different T2T genomes were built by different subsets of data types from HiFi, ONT UL, Hi-C, BioNano and strand-seq, ONT UL was required in most situations, without which a certain number of long repetitive regions with few detectable variations cannot be spanned, leading to assembly with large gaps^11, 14, 15^. However, due to the high price of ONT UL sequencing, it would be too expensive to do it for a population of genomes. Instead of ONT UL data, in most existing population-level sequencing projects (e.g., pan-genome), only Hi-C data and one of HiFi, CLR, or non-UL ONT data are available^16–18^. With the popularity of the Revio sequencer, the price of HiFi is reducing much faster than ONT UL and other types of sequencing, thus we believe “HiFi + Hi-C” mode will be the most popular choice for building larger and larger pangenome in the future. Third, compared with the complete haploid pangenome, building a haplotype-resolved complete diploid pangenome is even much more difficult. After all, it is still an unsolved problem for building a single diploid T2T genome. To conclude, in either a haploid or diploid way, with a widely-used assembly pipeline (e.g., hifiasm^7^ + 3dDNA^19^) for HiFi and Hi-C data and without a large amount of manual work, the assembled genomes in a pangenome may contain certain numbers of large gaps in repetitive regions (see **Fig. 1b** for an example in human centromeres).

**Fig. 1.**
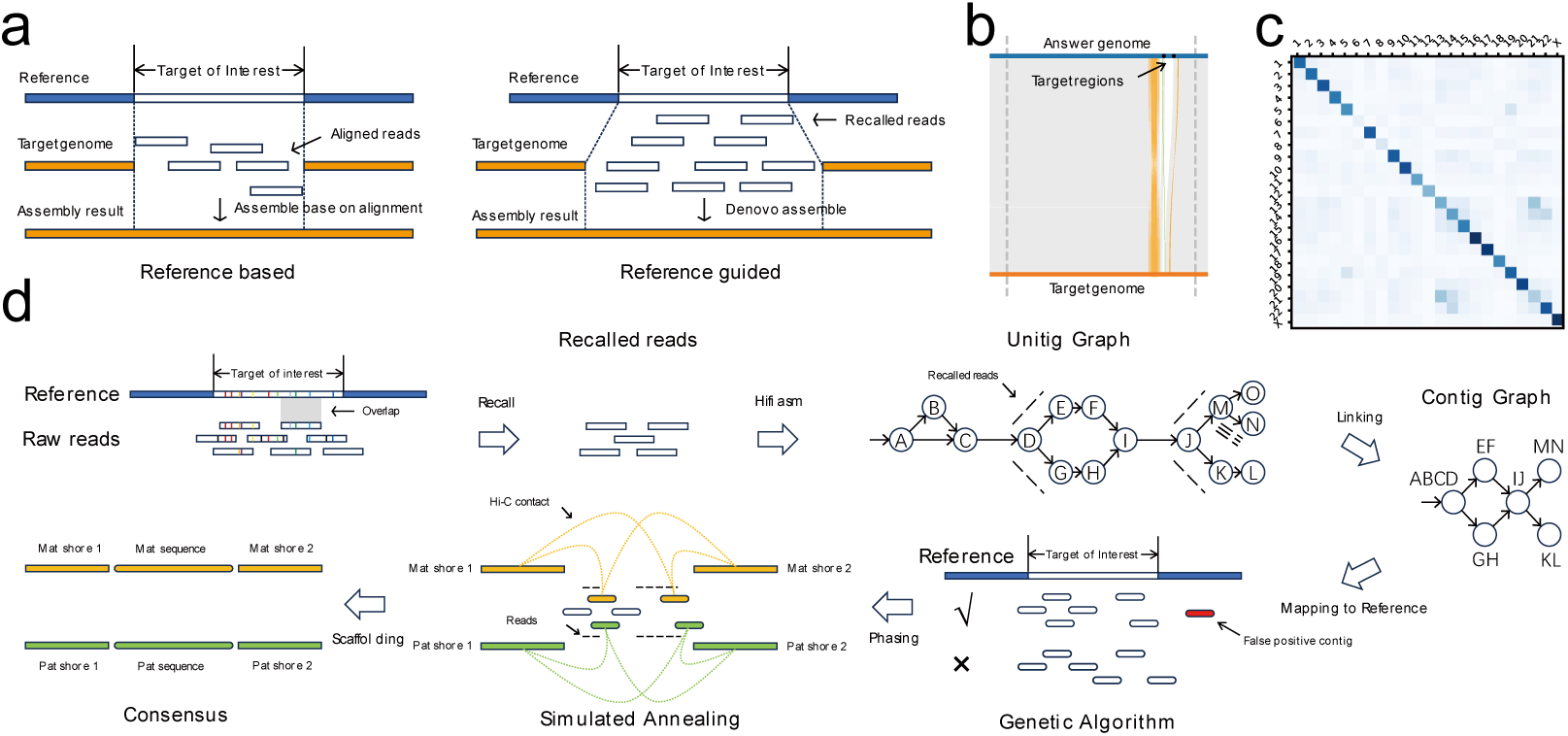
Overview of TRFill algorithm. **a**, Illustration of reference-based assembly and reference-guided assembly. **b**, A real gap illustration between 50Mb∼100Mb on chromosome 2 (paternal) of HG002 assembly generated by ‘hifiasm+3dDNA’ *de novo* pipeline using HiFi and Hi-C data. **c**, *k*-mer similarities between centromeric alpha satellite arrays on CHM13 and HG002 (maternal) chromosomes. **d**, Pipeline of TRFill algorithm.

To solve this problem and build complete pangenomes, an automatic algorithm is needed to achieve haplotype-resolved gap-filling in repetitive genomic regions in the chromosome-level assembles using only HiFi and Hi-C data. This paper reports an attempt to achieve this task on tandem repeats, which are usually much longer and thus more difficult to solve than interspersed repeats and other complex regions. Although this task looks extremely difficult in pure *de novo* genome assembly, it could be much easier when there exists a high-quality (T2T or near T2T) reference genome of closely related genome guiding the gap-filling of tandem repeats. Fortunately, for a species, this kind of high-quality single reference genome should be available before building a complete pangenome with a high possibility. Based on this idea, this paper proposed a new genome assembly mode called reference-guided genome assembly which is different from traditional *de novo* or reference-based assembly. More specifically, reference-guided genome assembly recalls the reads for each gap of interest by taking advantage of the corresponding regions in reference, and assembles these reads by comprehensively utilizing various information such as read overlaps, long-range links, and reference. As a proof-of-concept, we developed a reference-guided gap-filling algorithm called TRFill which is able to fill the gaps of tandem repeats in chromosome-level assembly in either haploid or diploid way using only HiFi and Hi-C data. Due to the difficulties of achieving this goal, a series of novel strategies were developed in TRFill for completing the steps such as accurate recall of reads, completely generating contigs from unitigs, determining contig positions and correct phasing.

The performance of TRFill was tested on alpha satellite repeats in human centromeres and the tandem repeats in tomato subtelomeres. On a diploid human genome (HG002), the assembly generated by a widely-used pipeline (hifiasm+3dDNA) of HiFi and Hi-C lost alpha satellite sequences in the vast majority of the centromeres. After the haplotype-resolved gap-filling with TRFill, the assemblies of alpha satellite sequences in 24 out of the 36 tested centromeres were improved in correctness, completeness or both. To test the performance on tomato genomes, we generated a dataset containing HiFi, ONT UL and Hi-C data for three tomato genomes. Guided by one of the tomato genomes built by all of HiFi, UL and Hi-C as a high-quality reference, the completeness and correctness of subtelomeric tandem repeats in 11 out of 15 and 15 out of 28 poorly-assembled subtelomeres were improved with TRFill in the other two tomato genomes (haploid) and a diploid simulated genome generated by merging them respectively, compared with the corresponding assemblies from standard “HiFi+Hi-C” pipeline. Furthermore, TRFill improved the completeness of the subtelomere for a recently published tomato pangenome with 29 genomes by 50% enabling a population-level analysis of the subtelomeric tandem repeats. With this analysis, the local law of sequence similarity of tandem repeats was found which in turn provides theoretical basis for reference-guided assembly. In addition, since the existing population-level analyses of tandem repeats mainly focus on centromeric regions^13, 20^, this study is a meaningful addition to non-centromeric regions in this area.

## Results

### Overview of TRFill algorithm

First of all, we introduce the concept of reference-guided assembly and its difference from the traditional reference-based assembly (**Fig. 1a**). As mentioned, reference-guided assembly recalls the reads for the genomic region of interest according to the reference and assembles them by comprehensively utilizing sequencing data and reference, which can be seen as a kind of combined method of the de novo one and the reference-based one. Although the reference-based and reference-guided assemblies both rely on the similarity between the reference genome and the genome to assemble, they have essential differences. Reference-based assembly assumes the ultra-high similarity between reference genome and the genome to assemble with differences only appearing at SNPs and small indels, requiring the vast majority of sequence in the reference to appear exactly once in the genome to assemble and in the same order. Therefore it performs the assembly directly according to the aligned position of reads on reference and generates inaccurate assembly when structural variations exist. Obviously, this assumption does not hold for tandem repetitive regions with a high possibility of structural variations (e.g., copy number variations). In contrast, the reference-guided assembly recalls the reads by only assuming the similarity in sequence composition, allowing the difference in the order and copy number of the sequence. Although the similarity in order is still needed to determine the order of the de novo assembled contigs according to the reference, the difference within the range of each contig is allowed.

On tandem repeats, the feasibility of the read-calling step process in reference-guided assembly is based on the assumption of local similarity regulation, i.e., the similarity in sequence composition (e.g., monomer similarity) between the corresponding region in reference and the genomic region to assemble is much higher than the similarity between the corresponding region in reference and any other regions in the genome to assemble (**Fig. 1c**). Otherwise, it is not possible to recall the reads for the gap to fill with both high precision and sensitivity. In this paper, we prove this assumption holds for at least two different kinds of tandem repeats: alpha satellite arrays in human centromeric regions and tomato subtelomeric tandem repeats (see the related sections for details).

Nevertheless, it is still a non-trivial task to completely recall the reads from the gap to fill and avoid recalling the false-positive reads from other tandem repetitive regions with similar sequence composition. To solve this problem, TRFill aligns the reads of the whole genome to the corresponding regions on the reference and generates a candidate set of reads with the ones from the gap region as completely as possible. To recall the reads from structural variation regions (regions in the genome to assemble but not in the reference) in the gap, we require only partial alignment rather than an alignment of the entire read by assuming the partial similarity of monomers between variation regions and reference. To remove the false-positive reads in the candidate set from other genomic tandem repeat regions, TRFill performs a hypothesis test checking if the representative *k*-mers in the aligned regions in the read are significantly different from the reference, by assuming a much higher similarity between monomers in the gap region than the one between a monomer in the gap and a monomer in any other tandem repeat region.

With the recalled reads, TRFill builds an unitig-level assembly graph (see **Supplementary Fig. 1** for a few examples) with hifiasm. It is noted that TRFill generates the contig-level assembly graph from unitig graph by itself instead of using the contig graph from hifiasm directly, of which the contigs may not be complete (see **Supplementary Fig. 2** for examples). TRFill traverses the unitig graph with breath-first-search (BFS) and depth-first-search (DFS)^21^, and pairs the in-edges and out-edges and removes the related wrong edges on each node (unitig) according to the numbers of supporting HiFi reads. Also, when meeting a long path and a single edge between a pair of unitigs both with enough supporting reads, TRFill prefers the long path to the single edge which proves to be the transitive edge with a high possibility by observation. Next TRFill removes the false-positive contigs assembled by wrongly-recalled reads and decides the order of the left contigs according to the alignments to the reference. It is non-trivial to finish these two tasks for two reasons: 1) the fragmented alignments due to the difference between contigs and the reference, and 2) multiple alignments of the same contigs on highly repetitive sequences. To solve these problems, a series of techniques were used in TRFill. First, for each contig, an alignment score for each possible alignment position of it on the reference is calculated by a dynamic programming algorithm^21^. The algorithm treats each aligned fragment on the contig and the corresponding fragment on the reference as an element and finds the longest aligned subsequence (similar to longest common subsequence, see Methods for formal definition) between contig and reference, the length of which is the alignment score. For each contig, a number of possible alignment positions with the highest alignment scores are selected as candidates, and then a global genetic algorithm^21^ is used to further select the appropriate alignment positions from the candidates for the contigs in a globally optimal way. The objective function of the genetic algorithm is to completely cover the corresponding gap region on the reference exactly one (haploid) or two (diploid) times, and each step of the algorithm tries to obtain a higher score of objective function than the last step by randomly removing one contig, adding back one contig, or changing the selected alignment position from candidates for one contig.

After deciding the positions of contigs on the reference, TRFill performs a phasing process for diploid genomes to assign the contigs to the different haplotypes according to the HiFi and Hi-C linkage information. More specifically, a simulated annealing algorithm^21^ is used with an objective function combining three sub-objectives: 1) maximizing the number of HiFi reads linking contigs from the same haplotypes, 2) maximizing the number of Hi-C read-pairs linking contigs and genomic regions closed to gaps (called shores) from the same haplotypes; 3) minimizing the length difference between the filled sequences of two haplotypes.

### Validation on Human Centromeric Alpha Satellite Arrays

We first validate the performance of TRFill algorithm on the alpha satellite sequences in the area of human centromeres. The alpha satellite sequences constitute approximately 3% of human genome and are composed of tandem repeats with repeat units (monomers) of ∼171 bp, multiple of which may form a higher-order repeat (HOR) unit of ∼2 kbp.^22^ For most species, the alpha satellite sequences are one of the longest continuous tandem repeats in genome.

In our experiments, the alpha satellite of a diploid human genome HG002 is chosen to be assembled with the human haploid T2T genome CHM13 as reference. Although the assembly of HG002 generated by the Human Pangenome Reference Consortium (HPRC) with a large amount of manual curation contains 195 gaps^23^, we still believe it is one of the most complete and correct assemblies existing for diploid genome and use it as the “ground truth”.

First, the whole-genome HiFi (36x) and Hi-C (69x) data of HG002 were assembled into a haplotype-resolved chromosome-level assembly following a widely-used *de novo* assembly pipeline (hifiasm-based contig assembly, 3D-DNA based scaffolding and Hi-C map based manual curation). Observe in **Supplementary Table 1** that there are in total only 3 of 36 centromeres with comparatively complete (completeness > 95%) alpha satellite regions. Among the 33 centromeres with incomplete alpha satellite sequences, there are 19 ones with completeness lower than 50% and 10 ones with completeness lower than 20%. On the other hand, the correctness of alpha satellite sequences is also poor. There are only 17 centromeres with the correctness of alpha satellite sequences > 90%, and 4 centromeres with ultra-low correctness (<20%).

Then, TRFill algorithm was run to reassemble the alpha satellite sequences for each centromere with CHM13 T2T assembly as reference. The position of each alpha satellite region on HG002 assembly was obtained by the synteny between HG002 assembly and CHM13 assembly with identified coordinates of alpha satellite sequences. Notably, we exclude acrocentric chromosomes (Chr13, Chr14, Chr15, Chr21, Chr22) from our experiments. In these chromosomes, the rDNA sequences are next to the centromeres, leading to the impossibility of identifying the exact starting or ending positions of alpha satellites regions in a poorly assembled area of HG002 assembly due to the high difficulty of assembling rDNA sequences, which may result in an inaccurate evaluation of TRFill algorithm. In the rest of the centromeres, observe that, for each of maternal and paternal haplotypes, the assemblies of alpha satellite sequences in two-thirds (12 out of 18) centromeres have been successfully improved (**Supplementary Table 2**). In terms of completeness, these 24 centromeres have been improved for 55% on average and 16 of them have achieved a completeness > 95% after reassembly. At the same time, the accuracy of all these alpha satellites have been either improved or kept at a very high level (almost 100% before and after reassembly) except for the one on chr18 (paternal) of which the accuracy reduced from 100% to 98.85% after reassembly (though the corresponding completeness was improved from 53.55% to 85.59%).

To further compare the assemblies before and after reassembly for these 24 alpha satellite sequences, we generated the synteny plots between them and the ‘ground truth’ (**Fig. 2a**, **Supplementary Fig. 3**). Observe that, although the assemblies generated by TRFill algorithm contains some imperfectness such as collapsed sequence (paternal chr20), false-positive sequences (paternal chr3) and sequence missing (maternal chr4), they are obviously better than the ones before reassembly. Notably, there are a few chromosomes such as chr1 and chr11, on which the alpha satellite sequences assembled on two different haplotypes are with significantly different lengths, but they both match the “ground truth” perfectly, although they are both guided by the same reference sequence. This phenomenon shows the advantage of reference-guided assembly proposed in this study over the traditional reference-based assembly, which should always generate assemblies with lengths same as the corresponding reference sequences.

**Fig. 2.**
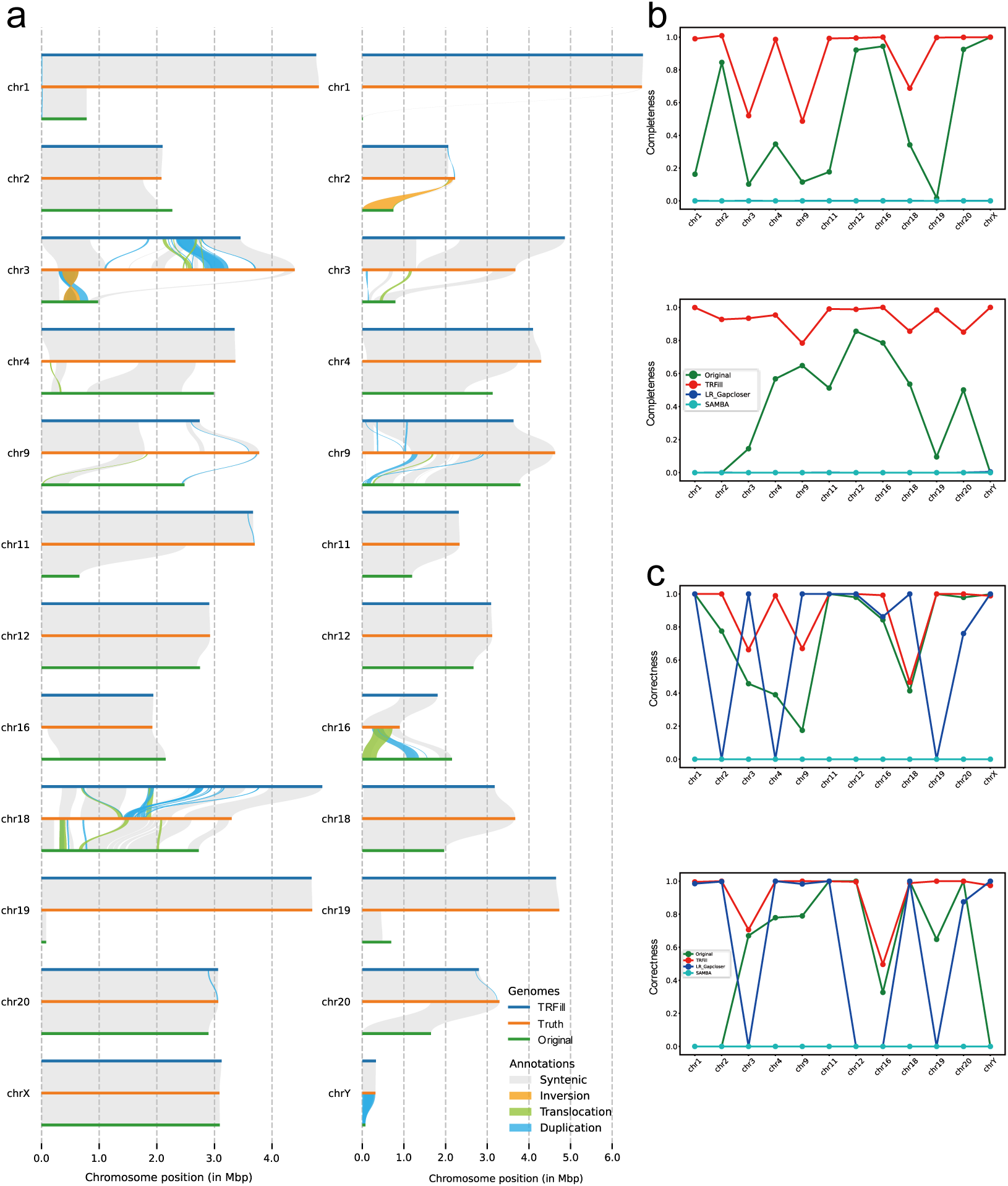
Validation on human centromeric alpha satellite sequences. **a**, Synteny plots generated by Syri between the original assemblies, the assemblies after TRFill reassembly of alpha satellite sequences and the “ground truth” assemblies for HG002 maternal (left) and paternal (right) chromosomes. Only the chromosomes with assemblies obviously improved by TRFill are shown. **b**, Completenss comparison of the original assemblies and the assemblies after gap-filling with TRFill, LR_Gapcloser and SAMBA on HG002 maternal (up) and paternal (bottom) chromosomes. **c**, Correctness comparison of the original assemblies and the assemblies after gap-filling with TRFill, LR_Gapcloser and SAMBA on HG002 maternal (up) and paternal (bottom) chromosomes.

Furthermore, we compare the assemblies from TRFill algorithm with the ones generated by two “state-of-the-art” *de novo* long-read based gap filling tools LR_Gapcloser^24^ and SAMBA^25^ (**Fig. 2b, 2c**). Observe that the completeness of their assemblies is almost close to 0 and even much lower than the assembly before reassembly. Although the accuracy is high (close to 100%) for some alpha satellite sequences, it is meaningless considering the ultra-low completeness. This result proves that the existing de novo gap filling tools are not useful for filling the gaps of long tandem repeats, because they all perform the progressive extension from two shores (the regions close to gap) which is very easy to be stopped on the tandem repeat with ultra-similar repeat units.

### Validation on Tomato Subtelomeric Tandem Repeats

Then we validated TRFill algorithm on subtelomeric tandem repeats of tomato genomes, which is a type of tomato-specific tandem repeats with repeat units of ∼181 bp (defined as *SolSTE181*). To perform this experiment, we built a dataset containing HiFi data of 38x, 40x and 44x, ONT UL of 48x, 37x and 40x, and Hi-C data of 130x for three homozygous tomato genomes Heinz1706 (*Solanum lycopersicum*), TS2 (*Solanum lycopersicum*), and TS281 (*Solanum lycopersicum var. cerasiforme*) respectively. The N50 of the three sets of ONT UL reads are 94,580 bp, 52,704 bp and 53,839 bp respectively. In this dataset, except for Hi-C data for Heinz1706 and HiFi data for all three genomes which were downloaded from the Sequence Read Archive (https://ncbi.nlm.nih.gov/sra) under BioProject PRJNA733299 and PRJNA756391^26^, other data was obtained in this study by sequencing. Besides this study, this dataset will be useful for testing other algorithms in genome assembly for future studies. For example, these data can be merged into the HiFi, ONT UL and Hi-C data for a simulated triploid genome and be used for testing the performance of polyploid genome assemblers.

In our experiment, first of all, the high-quality chromosome-level assemblies of these three genomes were generated with all of HiFi, ONT UL and Hi-C data by a widely-used pipeline (hifiasm-based contig assembly with HiFi and ONT UL data, 3dDNA-based scaffolding, and Hi-C map based manual curation). Among them, the high-quality assembly of Heinz1706 would be used as reference, while the ones of TS2 and TS281 would be used as “ground truth”. Observe that the subtelomeric tandem repeats of 19 subtelomeres have been successfully assembled in the high-quality assembly of Heinz1706 (**Supplementary Table 3**).

Second, we performed the validation on haploid genomes. We downloaded the assemblies of TS2 and TS281 generated by^26^ with hifiasm-based contig assembly, RagTag^27^ based scaffolding using a Heinz1706 assembly built by HiFi and Hi-C data as reference, and Hi-C map based manual curation. By comparing the subtelomeric tandem repeats in these two assemblies with their respective “ground truth” and with the high-quality reference genome (Heinz1706 assembly), observe that, there are 12 (TS2) and 3 (TS281) subtelomeres with comparatively low-quality (completeness < 98% or correctness < 98%) tandem repeats in these two assemblies respectively while the corresponding subtelomeres in reference contain high-quality tandem repeats (**Supplementary Table 4 haploid**). Thus, the reference genome was then used to guide the reassembly of the tandem repeats in these 15 subtelomeres. Observe that, after reassembly with TRFill algorithm, the tandem repeats of 11 out of the 15 subtelomeres were successfully improved in either one of completeness and correctness (the other one criteria not decreased) or both. Overall, for the 11 subtelomeres, the completeness and correctness were improved for 27% and 12% on average (**Fig. 3b, 3c, Supplementary Table 5 haploid**).

**Fig. 3.**
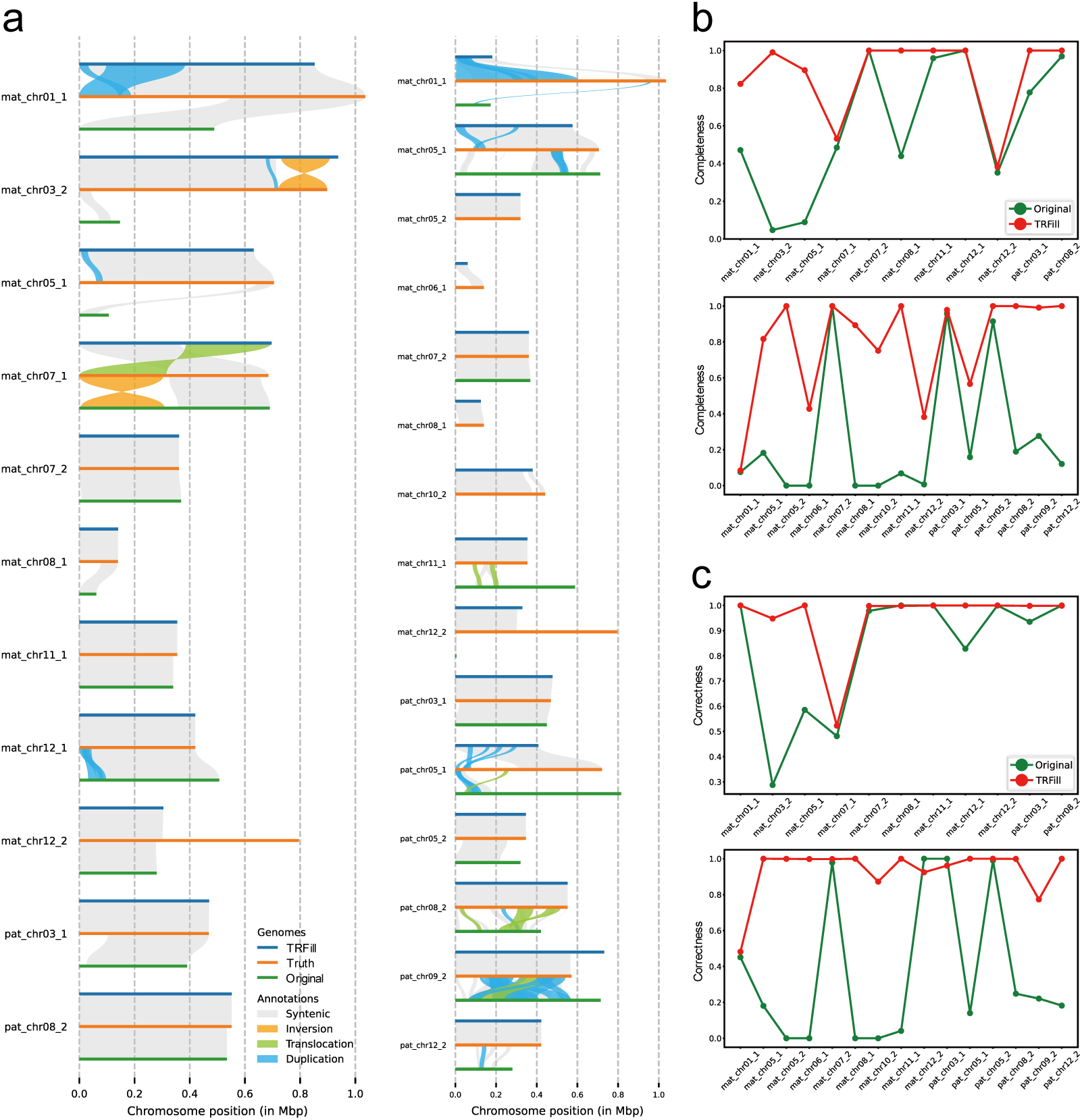
Validation on tomato subtelomeric tandem repeats. **a**, Synteny plots between the original assemblies, the assemblies after TRFill reassembly of subtelomeric tandem repeats and the “ground truth” assemblies for tomato haploid (left) and diploid (right) genomes. Only the chromosomes with assemblies obviously improved by TRFill are shown. In the figure, ‘mat’ refers to TS2 and ‘pat’ refers to TS281. **b**, Completenss comparison of the original assemblies and the assemblies after gap-filling with TRFill on tomato haploid (up) and diploid (bottom) genomes. **c**, Correctness comparison of the original assemblies and the assemblies after gap-filling with TRFill on tomato haploid (up) and diploid (bottom) genomes.

Third, we performed the validation on a simulated diploid genome with each of TS2 and TS281 as maternal and paternal haplotypes respectively. The HiFi and Hi-C data of it was generated by merging the respective data of TS2 and TS281, and was then assembled into a haplotype-resolved chromosome-level assembly using hifiasm-based contig assembly and phasing, 3dDNA-based scaffolding and Hi-C map based manual curation. Since the merged Hi-C data contains no inter-chromosomal signals between maternal and paternal chromosomes, simulated signals were generated to make up. In this assembly, there are in total 28 (17 on maternal and 11 on paternal) subtelomeres with comparatively low-quality (completeness < 98% or correctness < 98%) tandem repeats while the corresponding subtelomeres in reference contain high-quality tandem repeats (**Supplementary Table 4 diploid**). After reassembly with our TRFill algorithm, the tandem repeats in 15 of these 28 subtelomeres were improved in either one of completeness and correctness (the other one criteria not decreased) or both.

Overall, for the 15 subtelomeres, the completeness and correctness were improved for 53% and 57 % on average (**Fig. 3b**, **3c**, **Supplementary Table 5 diploid**). Finally, the synteny plots were drawn between the “ground truth” and the tandem repetitive regions before and after reassembly (**Fig. 3a**). Observe that, similar with the results on human datasets, intuitively the reassemblies are obviously better despite the existence of some structural variations with “ground truth” like duplications and inversions. Notably, the tandem repeats in some subtelomeres (e.g., ‘mat_chr05_2’ and ‘mat_chr08_1’ in diploid genome) were entirely missing before reassembly, while the corresponding ressemblies were in comparatively high completeness and correctness.

### Population-level analysis of *SolSTE181* sequences exhibits the local law of sequence similarity of tandem repeats

After validation of TRFill algorithm, we used it to reassemble the subtelomeric tandem repeats of 29 tomato genomes (including TS2 and TS281, **Supplementary Table 6**) in a pangenome with the 19 subtelomeric tandem repeats in Heinz1706 high-quality assembly as reference. Out of the 551 subtelomeres, TRFill algorithm generated longer tandem repeats than before for 494 ones, and the total length of tandem repeats were increased from 166,777,357 bp to 251,982,353 bp. The sequences were updated in the 494 subtelomeres, while the original sequences were kept in the other 48 subtelomeres (**Supplementary Table 7**). To validate the correctness of the assemblies, we compared the sequences of new assemblies with the original ones, and found that, in more than half of the 551 subtelomeres, the new assembly contains more than 99% sequence from the original one. Notably, in the new assemblies, there are sequences of in total 32,148,182 bp aligned to the contigs provided by^26^ which are not in their chromosome-level assemblies. Such phenomenon implies that TRFill has successfully recollects sequences missing from the original assemblies and put them inside.

With the updated subtelomeric tandem repeats, we performed a population-level analysis on the *SolSTE181* repeat units (monomer and HOR) by imitating a similar analysis on the alpha satellite array (*AthCEN178*) in *Arabidopsis thaliana* pan-centromere published recently^13^. First, the general analysis was performed on monomers. A monomer is defined as a ∼181-bp repeat unit, and 1,677,830 monomers were identified in total (**Fig. 4a**). Among these monomers, a high portion (149,012) appear uniquely in single genomes, and, same as *AthCEN178*, the monomers that are unique to single genomes tend to appear at high copy numbers (**Fig. 4d**). However, different from *AthCEN178*, the *SolSTE181* monomers shared by all or almost all genomes are also present at extremely high copy numbers. Furthermore, a consensus monomer was built from all monomers and the average variant frequency of each base was obtained (**Fig. 4b**). Different from *AthCEN178* monomer with an obviously higher variant frequency at the first half of bases than the second half, *SolSTE181* monomer is with a comparatively uniform distribution of variant frequency. In general, the average variation frequency of *SolSTE181* monomer is significantly higher than that of *AthCEN178*, indicating a lower stability of subtelomeres compared to centromeres. In addition, strong strand biases were observed among *SolSTE181* monomers (**Fig. 4c**).

**Fig. 4.**
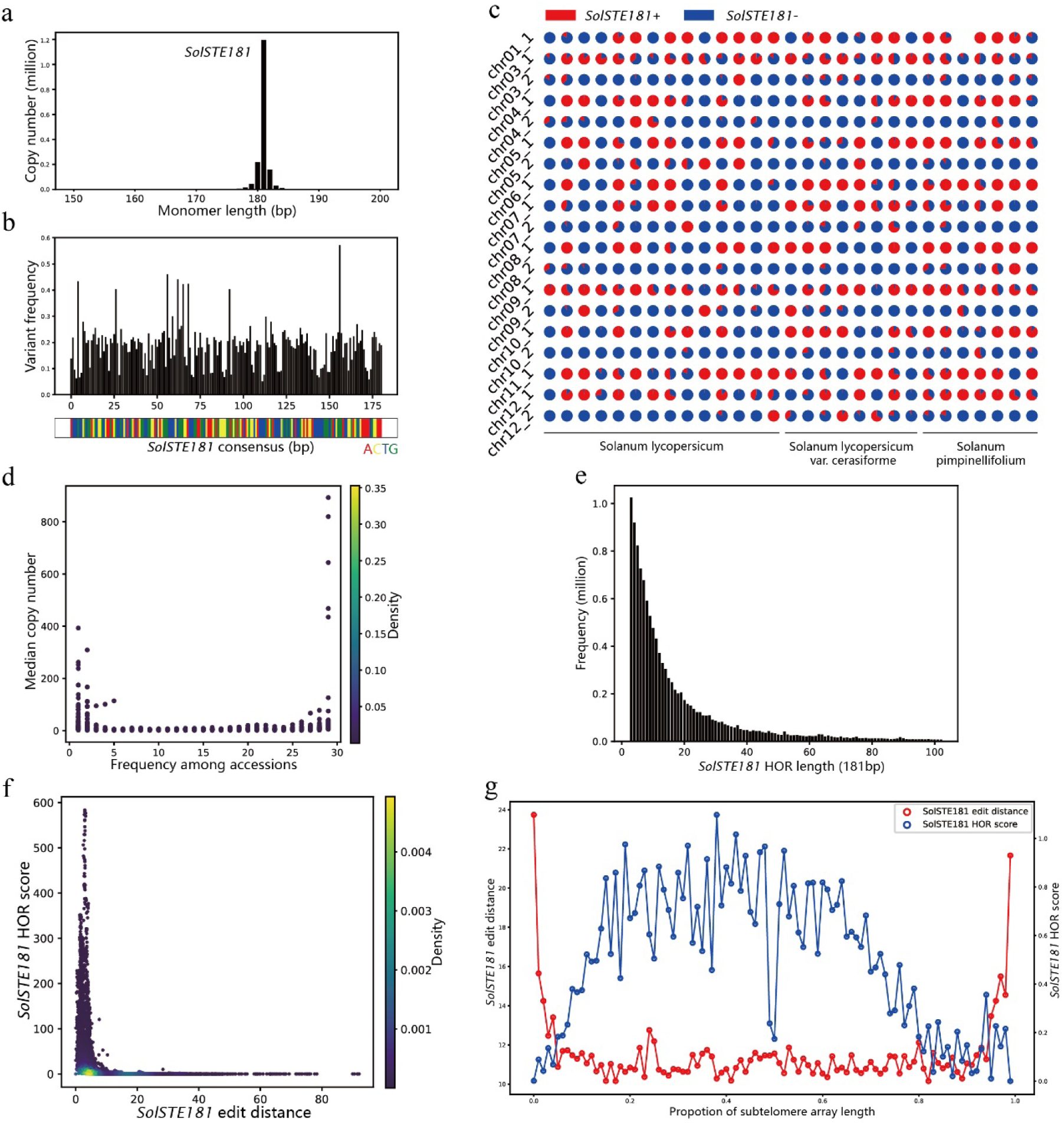
General population-level analysis on *SolSTE181* monomers and HORs. **a**, Length distribution of *SolSTE181* monomers. **b**, Variant frequency of each of the 181 nucleotides in *SolSTE181* concensus monomer, where the variant frequency is defined as ‘1 - the frequency of most frequent base’. The barcode DNA of the *SolSTE181* consensus is shown below (red: A; yellow: C; blue: T; green: G). **c**, Pie charts showing the proportions of *SolSTE181* monomers on the forward (red, +) and reverse (blue, -) strands respectively in 19 subtelomeres (rows) by 29 genomes (columns). **d**, Median SolSTE181 copy number per accession plotted against the number of accessions in which they were found. **e**, Length distribution of HORs with length described by the number of 181 bp monomers contained. **f**, Density plot of *SolSTE181* HOR scores versus edit distances from the *SolSTE181* consensus, across all accessions. **g**, Plot of *SolSTE181* HOR scores (blue) and edit distances (red) across the scaled length of all subtelomers.

Second, the general analysis was performed on HORs. In this study, HOR is defined as tandem repeat duplications of at least three consecutive monomers, each with no more than five substitution or single-base insertions/deletions per monomer pair. A total of 12,146,647 HORs of various lengths were found, showing a negative exponential length distribution (**Fig. 4e**), with the most frequent type being three-monomer HORs (∼543-bp), totaling 1,024,705 instances. To describe the repetitive nature of the HORs, the HOR score was defined for each monomer as the number of HORs to which the monomer belongs, normalized by the total number of monomers on the subtelomere it belongs. Furthermore, for each monomer, the edit distance from the consensus monomer of the subtelomere it belongs was calculated. Across the subtelomeres, the regions that represent 0.1 to 0.8 of the subtelomere scaled length exhibit the highest HOR scores and the lowest edit distances (**Fig. 4g**), indicating that these regions are relatively stable and show the greatest extent of repetition along the subtelomeres. Despite most monomers possess modest values of HOR scores and edit distances, there are ones with rather high HOR scores or edit distances (**Fig. 4f**). These represent the most repetitive monomers or monomers that have undergone significant mutations. Additionally, the absence of monomers that score high in both metrics suggests the overall high sequence-similarity of subtelomeres within the population.

Third, a monomer-based analysis was performed on the *SolSTE181* sequence similarity. The sequence-similarity comparison was performed 1) between the intra-genome monomers and inter-genome monomers, 2) between intra-chromosome and inter-chromosome monomers in the same genomes, and 3) between intra-subtelomere and inter-subtelomere monomers in the same genomes. Although there were no significant differences exhibited in sequence similarity for 1) and 2), the intra-subtelomere monomers exhibited obviously much higher similarity than the inter-subtelomere ones (3)) (**Fig. 5a**). This phenomenon indicates that, for *SolSTE181* sequences, the similarity between monomers is not uniformly distributed and is strongly related to the locations of the monomers. More specifically, the *SolSTE181* monomers can be divided into non-overlapped local regions in which the local monomers are significantly more similar. The previous study showed similar finding for human centromere, for which each local region is a centromere^22^. We call this as “The local law of sequence similarity of tandem repeats”. One step further, to study the sequence similarity between the corresponding subtelomeres of different tomato genomes, we calculated the monomer similarity between all pairs of 551 subtelomers of all 29 tomato genomes. From **Fig. 5b**, observe that the corresponding subtelomeres of closely-related genomes exhibited obviously higher sequence similarity than the different subtelomeres of even the same genomes. And the sequence similarity between corresponding subtelomeres is not much related to the evolutionary distance between genomes (**Fig. 5c**). This phenomenon indicates that “local region” could be a concept for a population of closely-related genomes rather than a concept for single genomes, and “The local law of sequence similarity of tandem repeats” may hold at population-level too. Since **Fig. 1c** shows a similar finding obtained on the centromeric alpha satellite sequences of human CHM13 and HG002 genomes, “The local law of sequence similarity of tandem repeats” may be a general law for tandem repeats (although need proof on more types of tandem repeats and more species), which will provide an important rationale for the reference-guided genome assembly proposed in this study of which the feasibility relies on the reference-based read-recalling.

**Fig. 5.**
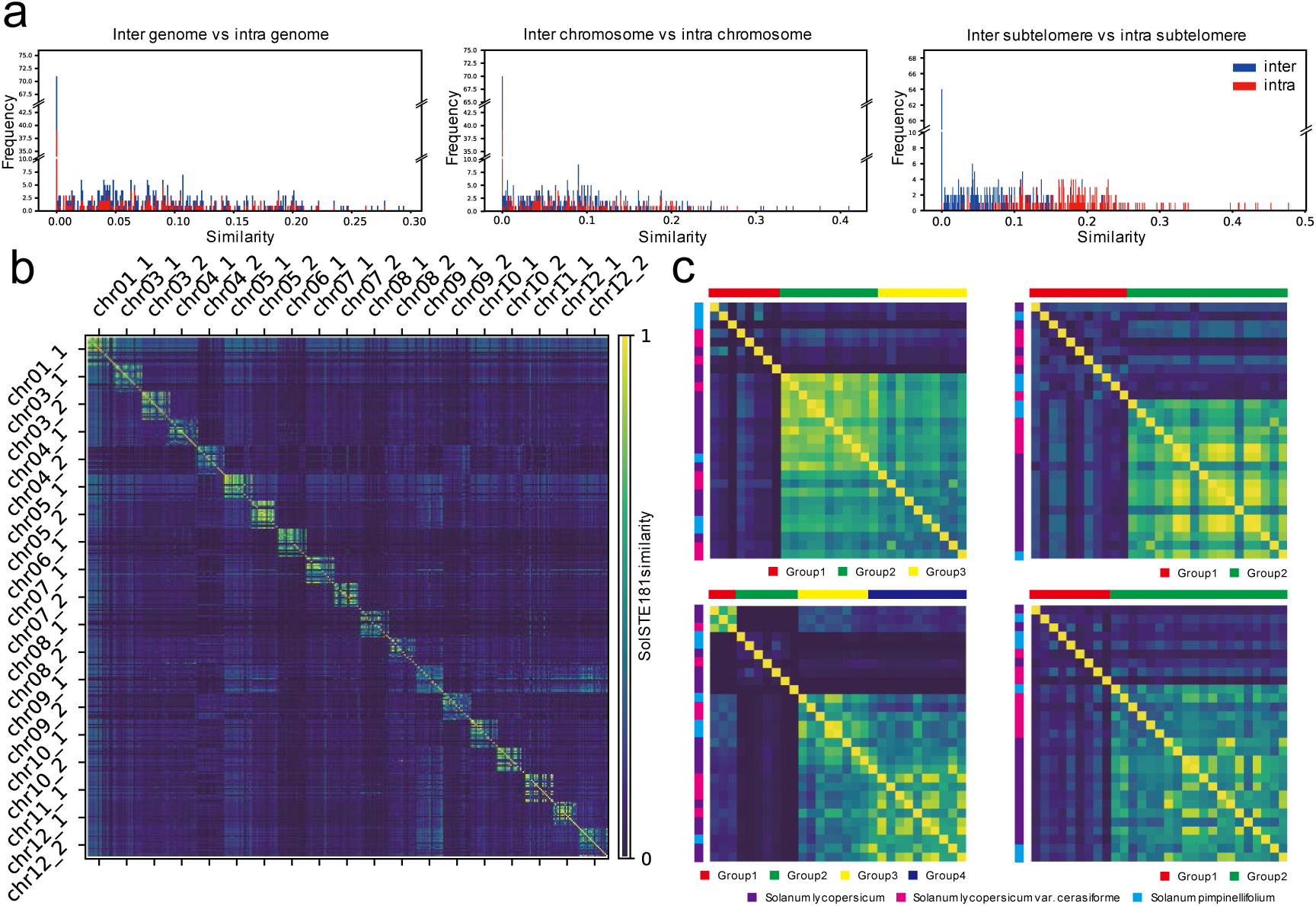
Monomer-based population-level analysis of *SolSTE181* sequence similarity. **a**, Comparision of similarity distributions of *SolSTE181* sequences between inter-(blue) and intra-(red) genome, chromosome and subtelomere respectively. **b**, Sequence-similarity between all pairs of *SolSTE181* sequences in 551 subtelomeres, with the subtelomeres at the same positions on different genomes put together. **c**, Each of the four matrices is generated from a submatrix on the diagonal of the matrix in **b** which contains the sequence-similarity information between 29 subtelomeres at a same position on 29 accessions. The generation process is to shift the order of accessions to ensure the accessions of which the corresponding subtelomeres in the same groups are put together. The groups are generated by hierarchically clustering subtelomeres according to the sequence similarity using the Python function fcluster from the package scipy. Different colors on X axes and Y axes represent the similarty groups and the three taxnomic groups respectively.

## Discussion

To generate population-level genomic map, the traditional method is to align the short reads of the population to the high-quality reference and obtain the assemblies according to the aligned positions on reference (called resequencing). Due to the inability of this method for correctly assembling sequences in structural variation regions, in recently years, large-scale pan-genomes were built for at least dozens of eukaryotic species in pure de novo assembly way. Without using any information from reference, the de novo method needs multiple types of high-depth sequencing data and a large amount of workload for manual curation in the assembly of each genome, especially when assembling long tandem repetitive genomic regions like centromere, ribosomal DNA and telomeres. It should not be questioned that it is a waste of resources to exploit a pure de novo method to generate population-level assemblies when one or more ultra-high-quality (e.g., telomere-to-telomere level) reference genomes for the same species are available. Therefore, it is needed to propose a genome assembly method which is able to take advantage of reference information to guide the assembly of new-sequenced genomes with comparatively low-priced long reads, and generate assembly results close to the ones from pure de novo method, which can be seen as a upgraded resequencing. In this study, we provide a solution called “reference-guided” genome assembly, which recalls the reads for the to-assemble genomic regions and assemble them in a de novo-like way. We implemented this idea into a proof-of-concept algorithm TRFill which is able to significantly improve the completeness and correctness of tandem repeats such as human centromeric alpha satellite arrays and tomato subtelomeric tandem repeats in both haploid and diploid way using comparatively low-priced sequencing data (e.g., HiFi and Hi-C) automatically.

Theoretically, the reference-guided assembly assumes the similarity between to-assemble genomic region and the corresponding region on reference be significantly higher than the similarity between non-corresponding regions. This assumption may hold with no doubt for non-repetitive regions, but for repetitive regions, it is not for sure due to the high similarity between the repeat units appearing in different genomic regions of some specific types of repeats (e.g. centromeric alpha satellite arrays, rDNA sequences, telomeres and transposons). Although, we proved the correction of this assumption on human alpha satellite arrays and tomato subtelomeric tandem repeats in this study, more studies are needed in the future on more species and more types of tandem and interspersed repeats such as rDNA, telomere and transposons to determine the application scope and limitation of reference-guided assembly. In addition, although the existing de novo assembly pipelines of long reads are already able to assemble non-repetitive regions completely and correctly, and this study focus on tandem repeats, it could be worthy to try reference-guided method for achieving complete and correct assembly of non-repetitive regions with ultra-low-depth long reads in the future.

Accurate process of read recalling is the cornerstone of reference-guided assembly. In TRFill algorithm, we use a parameter *δ* to describe the relationship between the expected rare k-mer number *μ* in the aligned read region and the observed rare *k*-mer number *y* in the aligned reference region as *μ* = *δy*(0 ≤ *δ* ≤ 1). Although *δ* is assumed to be globally constant in this study and the constant value estimated from Arabidopsis thaliana data works well for human centromeres and tomato subtelomeres, theoretically choosing different value for each gap could be more appropriate and may further improve the correction of read recalling process, considering that the difference in sequence composition between reference and query and the similarity in sequence composition within the related local region can be both different for different genomes and genomic regions. A potential method for more accurately determining *δ* value for each gap is to take advantage of the tandem repeat sequences next to the gap which has been correctly assembled before filling for estimation. Similarly, the value of *σ*^2^ representing the HiFi sequencing error rate can be more accurately estimated for each genome by taking advantage of the correctly assembled sequences in non-repetitive regions.

Although TRFill significantly improved the assembly quality on human and tomato data, we believe there is still a lot of room for improvement in the specific technical details. In fact, as a proof-of-concept algorithm, TRFill is designed only for proving the feasibility of reference-guided assembly on tandem repeats rather than providing a practical tool for users. We are confident that, after integrating the advanced techniques used in the ‘state-of-the-art’ long read assemblers in the future studies, the correctness, completeness, speed and space efficiency will all be significantly improved. For example, the unitig graphs (**Supplementary Fig. 1c**) of the centromeric alpha satellites failed to assemble by TRFill are mostly continuous and consistent with the “ground truth” in most regions, providing an opportunity for high-quality assembly of these sequences with improved algorithms in the future.

In TRFill algorithm, reference genome is used together with HiFi and Hi-C data for determining the order of contigs in scaffolding process. On one hand, it is another way of taking advantage of reference information besides the reference-guided read recalling and should be helpful for the contiguity of the regions with few HiFi and Hi-C links, while on the other hand, it can be seen as a disadvantage of TRFill algorithm because the correctness of assembly may be affected when existing structural variations larger than contig size though this situation should be rare for the long contigs from long reads. To address this tradeoff between correctness and contiguity, it could be worthy to try in the future an adaptive strategy which can adjust the weight of reference information according to the number and reliability of HiFi and Hi-C links. Also more studies are needed to explore an appropriate extent to which reference information should be utilized and how to use it.

Although TRFill algorithm improved the completeness and correctness of most tandem repeats in the experiments, there exist a small number of examples in which the new assemblies were worse than original ones. Therefore, in practical application, it is important to detect these regions for which the original sequences should be kept. Although there are many methods for evaluating the correctness of repetitive regions including web-experiments based methods like FISH and bioinformatic methods such as read coverage checking, they are all with too much workload for populational data and not easy to be carried out by people with limited bioinformatic techniques. Therefore, it is necessary to study if the existing automatic assembly evaluation tools are able to detect these poorly assembled genomic regions by TRFill and propose new evaluation tools.

To conclude, although our algorithm is still imperfect and can be improved in different ways, we believe it is a good start and a solid step towards the goal of complete populational genomics.

## Methods

For simplicity, in this section, we call the complete genome guiding the assembly process and the genome to assemble as reference and query genomes respectively. The gap region to assemble on query is called target area of interest (TOI) on query. Similarly, the corresponding region on reference genome is called TOI on reference. The input of the proposed TRFill algorithm contains: 1) the complete reference genome, 2) the chromosome-level query genome assembled by existing “HiFi+Hi-C” assembly pipeline (e.g., hifiasm + 3dDNA), 3) the whole genome HiFi and Hi-C reads for query and 4) the starting and ending positions of TOI on query. The output is the assembled sequence for TOI on query. More specifically, TRFill algorithm is composed of four phases: 1) recalling the reads for TOI on query according to TOI on reference; 2) assembling the recalled reads into contigs in de novo way; 3) determine the contig positions and orientations according to reference; and 4) phasing. The pipeline of TRFill algorithm is shown in **Fig. 1d**.

### TRFill Algorithm: Read Recalling

First of all, TOI region on reference is identified according to the synteny between reference and query obtained by Syri (v1.6)^28^. More specifically, the minimum-length continuous region on reference that covers the TOI region on query is selected as the TOI on reference. The whole-genome HiFi reads are then aligned to TOI on reference with winnomap2 (v2.03)^29^ with **‘**-x map-pb**’** option, and the reads with any one or more sub-regions aligned are selected into a candidate set. Considering the difference between reference and query, it is not required to have the entire read aligned.

Then a hypothesis test is performed to remove the false-positive reads (reads from other genomic regions) from the candidate set according to the matching of rare *k*-mers (in this study, we use 21-mers appearing less than 4 times in the whole reference genome). To obtain these rare *k*-mers, Jellyfish (v2.3.0)^30^ is run to count the appearing of the *k*-mers on reference and the frequent ones are removed. For each read in candidate set, we focus on its alignment with TOI on reference, and check if the rare *k*-mers appearing in the aligned reference region also appear completely in the aligned query region. More formally, for an alignment *A* between the aligned region *R* on reference and the aligned region *Q* on read, let *Y* and *X* (*X* ≤ *Y*) be the number of rare k-mers appearing in *R* and the number of rare k-mers appearing in both *R* and *Q* respectively. Intuitively, the test is to check how close are the value (*x*) of *X* to the value (*y*) of *Y*. We assume *X* to be normal distributed (*X*∼*σ*(*μ*, *σ*^2^)) with *μ* representing the number of rare *k*-mers appearing in both *R* and the corresponding region in the real query genome, and *σ*^2^ describing the sequencing error rate. Since the real query genome and *μ* are unknown, we further assume *μ* can be a function of *y* if *R* and *Q* are homologous regions (i.e., *R* and *Q* are from the same local tandem repeat regions of different genomes such as the centromeres of the same chromosome in two different genomes). Considering *μ* is always smaller than or equal to *y* according to their definition, we assume the function as *μ* = *δy*(0 ≤ *δ* ≤ 1). Although theoretically *δ* is affected by many factors such as the difference in sequence composition between reference and query and the similarity in sequence composition within the related local region of tandem repeat for both reference and query, for simplicity, it is assumed to be constant in this study. Since *σ*^2^ is only determined by the sequencing technology and only HiFi reads need to be recalled in this study, *σ*^2^ is also assumed to be constant. With these assumptions, the probabilistic density function of *X* can be described as

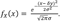

Notably, *δ* and *σ*^2^ are two important parameters of TRFill algorithm which are strongly related to the performance. More discussion about them is shown in the Discussion section. In this study, the values of *δ* and *σ*^2^ were estimated on centromeric regions of Arabidopsis thaliana. ColCEN, a near-T2T-level genome of Arabidopsis thaliana (accession Col-0) with almost complete centromeric regions^31^, was used as reference and the HiFi reads from the centromeric regions of another genome (accession Ler-0^13^) of Arabidopsis thaliana were aligned to the corresponding centromeric sequences of ColCEN. To obtain the HiFi reads from each centromere of accession Ler-0, we align the whole-genome reads to its assembly containing comparatively complete centromere, and keep the reads with high-quality alignments. In this way, we obtained a set of alignments (*A*_*i*_, 1 ≤ *i* ≤ *σ*) between HiFi reads from the centromeric regions of accession Ler-0 and the centromeric regions of

ColCEN. For each *A*_*i*_, the related variables *R*_*i*_, *Q*_*i*_, *X*_*i*_, *Y*_*i*_, *x*_*i*_, *y*_*i*_ can be defined same as *R*, *Q*, *X*, *Y*, *x*, *y* respectively. Then the probabilistic density function of *X*_*i*_ can be described as

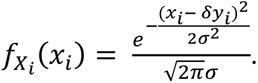

By combining the probabilistic density functions of all *X*_*i*_ (1 ≤ *i* ≤ *N*), the log likelihood function is obtained as

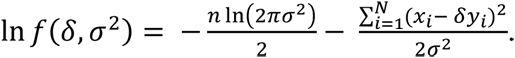

By maximize the log likelihood function, we have

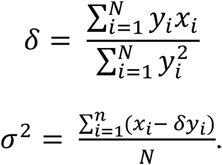

With the estimated *δ* and *σ*^2^, a two-sided test is performed on each alignment *A* (with *R*, *Q*, *X*, *Y*, *x*, *y* defined above). Let 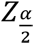 be the value of standard normal distribution at a certain significance level (e.g., *α* = 0.05). If *x* falls into 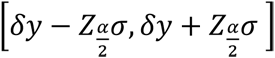, we recall the read. Otherwise, it is seen as a false positive read and discarded. For a read with multiple alignments to TOI on reference, we perform the test on all alignments and recall this read as long as one alignment passes the test.

### TRFill Algorithm: Contig Assembly

After recalling HiFi reads for TOI of query genome, we assemble them into contigs in *de novo* way. First of all, a unitig-level assembly graph is built from these recalled reads with hifiasm (v0.19.5-r587). The default settings are used for diploid genome and **‘**-l0**’** option is used for haploid genome. Notably, although hifiasm also provides contig-level assembly graph, examples show that the contigs assembled directly from hifiasm are with low completeness (**Supplementary Fig. 2**). Thus, TRFill algorithm generates contigs from unitig graph by itself.

To achieve this, the HiFi reads are aligned back to the unitig graph by winnowmap2 (v2.03) and the number of supporting reads for each edge is counted. Subsequently, Breadth First Search (BFS) is used to traversed the unitig graph from zero-indegree nodes, and generate a simplified graph which contains the same vertices as unitig graph and all the edges in unitig graph traversed in BFS. During the traversal, at each branching node v, the following rules are followed. (1) Different from the traditional BFS allowing each node visited only once, we allow v to be visited for ⌊(*id* + *od*) / 2⌋ times in the whole BFS traversal, where *id* and *od* are the indegree and outdegree of *ν*. (2) For a successor node *w* of *ν*, if there exists a longer path from *ν* to *w* in which *o* is the direct successor node of *ν*, *o* is chosen for visiting next rather than *w* for this time of visit of *ν* (i.e., edge *ν* → *w* is seen as a transitive edge) (see *A*, *B*, *C* in unitig graph of **Fig. 1d** for an example). Notably, there is still chance for *w* to be visited from *ν* in the next time of v being visited. (3) If the situation of (2) does not exist, among all the successors of *ν* with enough supporting reads, the one with the largest number of supporting reads is chosen for visiting for this time of visit of *ν* (see *M*, *O*, *N* in unitig graph of **Fig. 1d** for an example). Similarly, there is still chance for other successors to be visited from *ν* in the next time of *ν* being visited. (4) If no successor of *ν* has enough supporting reads, all successors of *ν* are chosen to visit for this time of visit of *ν* (see *D*, *E*, *G* in unitig graph of **Fig. 1d** for an example). After finishing the BFS traversal and building the simplified graph, the order of the nodes visited and the corresponding timestamps are kept. A node visited for more than one times in BFS appears for multiple times in the order.

Next, Depth First Search (DFS) is used to traverse the simplified graph to generate contigs. Same as BFS, in the traversal of DFS, each node is allowed to be visited for ⌊(*i* + *di*) / 2⌋ times where *id* and *od* are the indegree and outdegree. At each branching node *ν*, among all the successors of *ν* with the maximum number of visits not reached, the algorithm chooses the one with earliest timestamp in the BFS traversal to visit. At the same time, the algorithm records that the graph needs to be cut at *ν* after finishing the traversal to generate discontiguous contigs (the branches left in the simplified graph are seen as unsolvable). However, there is a special situation (loop) where the related branches can be solved. **Supplementary Fig. 4a** shows an example, in which *E* → *A* → *B* → *C* → *A* → *D* is a unique path with no ambiguity and is able to generate a continuous contig, although *A* is a branching node. To distinguish this special situation from other unsolvable branching nodes, in the algorithm, as long as a node is revisited and a loop is generated, the record of cutting at it is removed. In this example, although *A* is recorded to be cut after finishing traversal at the first time of its visit, the record is removed at the second time of its visit and thus *E* → *A* → *B* → *C* → *A* → *D* is still a continuous contig. In some situations where the unsolvable branching node is nested in a solvable loop. **Supplementary Fig. 4b** shows an example in which *B* is an unsolvable branching node and will be cut after traversal while the other part of the loop is solvable. Thus, finally, three contigs will be generated as *E* → *A* → *B*, *C* → *A* → *D* and *F*.

After DFS traversal, the simplified graph is cut at the branching nodes recorded by following a few rules. Let *ν* be a branching node recorded. If *ν* has one precursor and multiple successors, the outgoing edges of *ν* are cut and the incoming edge is kept. If *ν* has multiple precursors and one successor, the incoming edges of *ν* are cut and the outgoing edge is kept. If *ν* has multiple precursors and multiple successors, all the incoming edges and outgoing edges of *ν* are cut.

### TRFill Algorithm: Determining Positions and Orientations of Contigs

After obtaining the contigs, we determine the contig positions and orientations according to the reference. First of all, the contigs are aligned to the TOI of reference with winnowmap2 (v2.03) and a filtering process is done. In the filtering process, the contigs satisfying any one of the following criteria are seen as false-positive contigs assembled by recalled false-positive reads and removed: (1) the contigs that do not have an alignment with TOI of reference with length longer than 10% of their own lengths; (2) the contigs of which the total length of alignments to TOI of reference is not more than twice the total length of alignments to other genomic regions on reference.

After filtering, we decide the positions and orientations of the left contigs according to their alignments to the reference. This process is non-trivial due to the fragmented alignments caused by the differences between reference and query genomes and the multiple alignments caused by the highly repetitive sequences.

To solve the problem of fragmented alignments, we designed an optimization model. Assuming a contig is aligned to TOI of reference and a set of fragmented alignments are obtained. For each alignment, we define the aligned regions on reference and contig both as elements. By collecting the elements on the contig of all these alignments and sorting them by their starting positions on the contig, a sequence

*L*_*c*_ [1 … *N*][1 … *N*] of elements on contig side is obtained. On reference side, we try all sub-region of TOI with length equal to the contig length and starting from an element aligned to *L*_*c*_[1]. For each of these sub-regions, in the similar way as the contig side, a sequence *L*_*r*_ [1 … *M*] of elements on reference side is obtained. Then the alignment score between contig and this sub-region can be calculated by the length of the Longest Matching Subsequence (LMS, see Definition 1 for details) between *L*_*c*_and *L*_*r*_where the ‘matching’ in LMS represents the ‘alignment’ between elements.

### Definition 1 (Longest Matching Subsequence)

Input: A sequence *A*[1 … *N*] of elements, a sequence *B*[1 … *M*] of elements, and a matching function *f* in which *f*(*x*, *y*) = *True* if element *x* in *A* and element *y* in *B* match. Output: a subsequence *A*’ of *A* and a subsequence *B*’ of *B* satisfying (i) the length |*A*’| of *A*’ is the same as the length |*B*’| of *B*’ and (ii) for any position *k* (1 ≤ *k* ≤ |*A*’|), *f*(*x*, *y*) = *True* for the element *x* from *A*’ and element *y* from *B*’ both at kth position.

Since the problem of LMS is very similar with the classical problem of Longest Common Subsequence (LCS)^32^, the time complexity of the optimal algorithm is also *O*(*NM*) which may not be acceptable for some contig with very fragmented alignments.

To speed up, in practice, we obtain an approximate solution of LMS by solving another classical problem called Longest Increasing Subsequence (LIS)^33^. For the sequence *Lr* of elements on reference side, a corresponding coordinate sequence *L*[1 … *M*] of same length is built according to the sequence *Lc* of elements on contig side. Formally, for any *j* (1 ≤ *j* ≤ *M*), *L*[*j*] = *i* if *Lr*[*j*] is aligned to *Lc*[*i*]. Then a longest increasing subsequence *LL*[1 … *P*] satisfying the following criteria is obtained for *L* with a binary search algorithm^34^ of time complexity *O*(*N* + *MlogM*) : 1) *LL* is a subsequence of *L*; 2) for any *p*, *q* (1 ≤ *p* < *q* ≤ *P*), *LL*[*p*] ≤ *LL*[*q*]; 3) there exists no other subsequence of *L* satisfying 1) and 2), but with length larger than the length of *LL*. The length of the longest increasing subsequence is used as the alignment score between the contig and this subregion of TOI on reference. For each contig, we keep at most seven subregions with the highest alignment score as candidate alignment positions.

Next, a genetic algorithm is used to select a subset of these contigs and determine the position of each selected contig from its candidate alignment positions in an optimal way. In the genetic algorithm, each contig, each alignment postition and each feasible combination of candidate alignment positions of contigs are considered as a ‘locus’, an ‘allele’, and an ‘individual’ respectively (‘locus’, an ‘allele’, and an ‘individual’ are all terms in the discourse system of genetic algorithm). By simulating the evolution process involving ‘switching’ (switching alignment position of one contig between two ‘individuals’), ‘variation’ (delete or insert one contig from or to ‘individual’) and ‘natural selection’ (select ‘individuals’ with higher score in objective function to next iteration with higher probability), we search for the optimal ‘individual’ that maximizes the objective function (also called the ‘adaptivity’ of ‘individual’ in the discourse system of genetic algorithm). The objective function of optimization comprises of two parts, the coverage related part and the distance related part. The coverage part aims to ensure that the contigs collectively cover every part of the TOI of reference exactly *r* times, where *r* represents the ploidy of the query genome (e.g., *r* = 2 for diploid genome). The distance part aims to maximize the distance between contigs to reduce the extent of overlap between them. Notably, these two objectives can be contradictory: covering the reference more extensively tends to bring contigs closer together.

Formally, let *l*_*t*_ be the total length of the subregions of TOI on reference possessing a coverage of *t*, and *d*_*ij*_ (*i*, *j* ∈ [1, *C*]) be the distance between alignment midpoints of contig *i* and contig *j* on the TOI of reference where *C* represents the total number of contigs. The objective functions for haploid and diploid genomes are shown as follow.

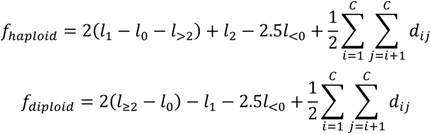

Notably, *l*_<0_ represents the total length of a special type of subregions with coverage 0. For each of this type of continuous subregions, the contigs aligned left and right of it belong to the same connected component in contig-level assembly graph. Since theoretically this kind of situation should not appear, we distinguish it from other 0-coverage subregions and give it a higher penalty. *M*_0_ represents the total length of the rest subregions with coverage 0.

The genetic algorithm proceeds in the following steps with default parameters:

1. Initialization: Randomly generate 500 ‘individuals’.
2. Termination condition: Calculate the ‘adaptivity’ (score of objective function) for each ‘individual’. The algorithm terminates if either of the following criteria is satisfied: 1) the 10 ‘individuals’ with the highest ‘adaptivity’ have the same ‘adaptivity’; 2) the algorithm has been executed for 200 iterations.
3. Natural selection: Randomly select 60 ‘individuals’ from the 500 ones according to their ‘adaptivity’ (the ‘individuals’ with higher ‘adaptivity’ can be selected with higher probability).
4. Switching: Randomly select two ‘individuals’ from the 60 ones, and use them to generate two ‘offspring individuals’ by randomly ‘switching’ their ‘alleles’ (the alignment position of the contig) at one same ‘locus’ (contig).
5. Variation: For each of the ‘offspring individual’, add a ‘mutation’ with a probability of 0.05 by either deleting one ‘locus’ (contig) and its corresponding ‘allele’ (the alignment position of the contig) or adding one ‘locus’ (contig) and one of its ‘allele’ (one of the candidate alignment positions of the contig).
6. Repeat (4)-(5) until 300 ‘offspring individuals’ are generated, and then return to (2) for next iteration.

After the termination of genetic algorithm, we select the subset of contigs and the combination of candidate alignment positions of contigs in the subset from the ‘individual’ with the highest ‘adaptivity’ as the final result.

### TRFill Algorithm: Phasing

After deciding the positions of recalled contigs on reference, the final step is to generate the sequence for TOI on query. For haploid genome, the contigs are directly connected into scaffolds by adding ‘N’s. For diploid genome, a phasing process with HiFi and Hi-C data and homology between contigs is performed before generating the scaffolds. Since HiFi and Hi-C are two kinds of linkage information with different resolution, we use them in a different way. HiFi reads are used to link the contigs in TOI belonging to the same haplotypes, while Hi-C signals are used to link the contigs and the genomic regions next to TOI in query genome (called shores). To obtain allele-specific Hi-C signals, we only consider Hi-C pairs with both ends on unique *k*-mers (*k*-mers appearing only once on contigs and shores, *k*=31). More specifically, Hi-C reads are positioned on contigs and shores according to the positions of unique k-mers contained rather than sequence alignment. However, when obtaining HiFi signals, due to the low-density of unique *k*-mers and the low-length of contigs in repetitive regions, the HiFi reads with both ends on unique *k*-mers on two different contigs are usually very rare and not enough for guiding phasing (it works for Hi-C because only one pair needs to be in repetitive region). Therefore, the HiFi signals are obtained by all HiFi reads aligned to two different contigs (by winnowmap2 (v2.03)), rather than requiring aligned on unique k-mers. Nevertheless, we assume that the HiFi signals linking the contigs of same haplotypes are still much more than the ones linking the contigs of different haplotypes. For homology, it is assumed that the contigs with higher homology should belong to the same haplotype with lower possibility. To calculate the homology between contigs, we set a homologous coefficient between each pair of contigs which is calculated by the number of common *k*-mers.

With the HiFi and Hi-C signals and homologous coefficients, we build an optimization model for phasing. Let *L*_*i*_ be the length of sequence *i* (contig or shore), *H̄* be the average value of the homologous coefficients of all pairs of contigs, *MAT_shore_*, *PAT_shore_*, *MAT_contig_*, *PAT_contig_*) be the sets of shores belonging to and the sets of contigs assigned to the maternal and paternal haplotypes respectively (the collapsed contigs belong to both *MAT_contig_* and *PAT_contig_*), *HIC_ij_,HIFI_ij_,D_ij_,H_ij_* be the number of Hi-C signals, the number of HiFi signals, the distance, the homologous coefficient between sequence *i* and sequence *j* respectively. The objective function is given by

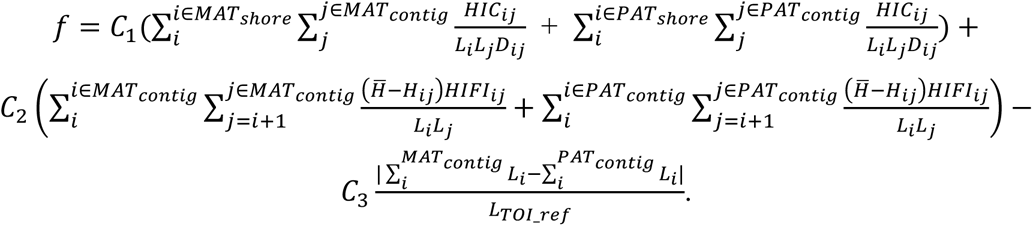

Intuitively, the objective function is composed of three parts. The first part is to maximize the Hi-C signals linking the shores and contigs in the same haplotypes. The second part is more complicated and should be considered in two situations. In the situation with a positive value of *H̄* − *H*_*ij*_ (low homology between contig *i* and *j*), *HIFI*_*ij*_ contributes positive to the objectively function representing the objective of maximizing the HiFi signals connecting the contigs in the same haplotypes. And at the same time, a higher value of *H*_*ij*_ leads to a lower *H̄ H*_*ij*_ which results in a lower score of objective function, representing the negative effect of homology. In the situation of a negative value of *H̄* − *H*_*ij*_ (high homology between contig *i* and *j*), *HIFI*_*ij*_ contributes negative to the objectively function. This arrangement is to deal with the homologous contigs from different haplotypes which may possess a large number of wrong-aligned HiFi reads connecting them due to the high sequence similarity. The third part is to minimize the length difference between the assembled maternal and paternal TOI sequences. Nevertheless, affected by the other two parts of the objective function, the algorithm is still able to generate accurate results for haplotypes with differential TOI lengths.

To maximize the objective function within an acceptable time, a simulated annealing algorithm is used. In this process, each contig can be in one of four states: 0 (discarded), 1 (maternal), 2 (paternal), or 3 (collapsed). The algorithm proceeds in the following steps with default parameters:

1. In initialization, for each contig, randomly set its state. And set the global maximum of objective function to be negative infinity.
2. Calculate the value of objective function. If this step has been executed for 1000 times, exit.
3. Randomly select a contig and change its state to a new one if doing so improves the value of objective function.
4. Repeat step (3) until the value of objective function remains unchanged for 100 consecutive iterations and thus a local maximum is obtained. If the local maximum surpasses the global maximum, set it as the global maximum.
5. Randomly changing the states of a percentage (50%) of contigs to new states for getting rid of local optimum. Return to step (2).

For either haploid or diploid assembly, the final assembly may contain some regions of shores due to some related false-positive reads recalled. To address this, TRFill trims such sequences according to the alignments of the scaffolds to the shores. Finally, the gaps of each haplotype are filled with the respective trimmed scaffolds.

### Validation on Human Centromeric Alpha Satellite Arrays

TRFill algorithm is tested on a diploid human genome HG002 (NA24385). The HiFi data (36x in coverage) and Hi-C data (69x in coverage) of HG002 are generated by the Human Reference Pangenome Consortium (HPRC)^23^. The haplotype-resolved assembly generated by the HPRC with a large amount of manual curation is used as “ground truth” to evaluate results of our algorithms.

To generate the chromosome-level diploid assembly of HG002 as the input of TRFill algorithm, hifiasm (v0.19.5-r587) with default parameters was used to generate haplotype-resolved contigs using HiFi and Hi-C data. Then the final assembly is generated by the haplotype-resolved scaffolding using 3D-DNA (branch 201008) with **‘**-r 0**’** option and manual curation based on Hi-C heap map with Juicebox^19^.

The human haploid T2T genome (CHM13 v2.0) is used as reference to guide our assembly of HG002. The positions of TOI regions on HG002 are decided according to the positions of centromere alpha satellite sequences on CHM13. The starting and ending positions of centromere alpha satellite sequence of each chromosome on CHM13 were determined by combining the downloaded annotations and results from Tandem Repeat Finder (TRF, v4.09.1)^35^. Notably, for each chromosome of CHM13, only the longest continuous alpha satellite sequence is used if this centromere contains multiple pieces of alpha satellite sequences. Then Syri (v1.6) with default parameters is used to build the synteny between CHM13 and two haplotypes of HG002, and the regions on HG002 aligned to the selected alpha satellite sequences are chosen as TOI regions. Notably, there may be a small proportion of genomic sequences within TOI regions assembled in the chromosome-level assembly of HG002. These regions are removed from the assembly and generating a continuous gap for each TOI. Then our TRFill algorithm, and two existing gap filling tools LR_Gapcloser and SAMBA (v4.1.0) are used to fill the gaps of TOI regions. TRFill algorithm was run with CHM13 assembly as reference. To evaluate the quality of assemblies in contiguity, completeness and accuracy, we used some criteria based on sequence alignment. To generate the alignment-based criteria, the assembly of TOI is aligned to the ground truth with winnowmap2, and the regions in assembly with continuous alignments longer than 100 kb are counted as correct genomic regions. The ratio of the total length of correct regions to the total length of ground truth is defined as the *completeness*, while the ratio of the total length of correct regions to the total length of assembly is defined as the *correctness*. Notably, since LR_Gapcloser and SAMBA are not able to generate continuous correct regions longer than 100 kb, we have to define the correct regions of their assembly as all aligned regions, which is beneficial for them. In addition, *LIS score* is defined as the ratio of the length of LIS between the unique *k*-mer sequences in “ground truth” and the assembly to the number of assembly unique *k*-mers (see section ‘TRFill Algorithm: Determining Positions and Orientations of Contigs’ in Methods for the generation of LIS).

### Validation on Tomato Subtelomeric Tandem Repeats

To test TRFill algorithm on tomato subtelomeric tandem repeats, we built a dataset including HiFi, ONT UL and Hi-C data of three tomato genomes (Heinz1706, TS2, and TS281) by doing ONT UL sequencing for all of them, Hi-C sequencing for TS2 and TS281, and downloading Hi-C data for Heinz1706 and HiFi data for all of them. More specifically, the young leaves were collected from an individual tree planted in the field of Agricultural Genomics Institute at Shenzhen, Chinese Academy of Agricultural Sciences (Guangdong province, China). In the ONT UL sequencing, high-molecular-weight genomic DNA selected with SageHLS HMW library system (Sage Science) and processed with the Ligation sequencing 1D Kit was used for the library construction using similar approach as described in^36^ and sequenced on the Nanopore PromethION platform at the Genome Center of Grandomics (Wuhan, China). In the Hi-C sequencing, the library was constructed and sequenced using similar approach as described in^37^ and sequenced using the Illumina NovaSeqX-plus platform at the Genome Center of Grandomics (Wuhan, China).

To build the reference genome (Heinz1706) and two “ground truth” genomes (TS2 and TS281), we assembled them separately in a do novo way. For each of them, hifiasm (v0.19.5-r587) with ‘-ul’ and ‘-l0’ options was used to generate contigs with HiFi and ONT UL reads, and 3D-DNA (branch 201008) with **‘**-r 0**’** option was used for scaffolding and generating chromosome-level assembly using Hi-C data.

In the experiments of haploid genome, the subtelomeric tandem repetitive regions of each of TS2 and TS281 genomes were used to assemble with TRFill algorithms guided by Heinz1706 reference genome. First, the chromosome-level assembly of TS2 and TS281 was downloaded from the SolOmics database (http://solomics.agis.org.cn/tomato/ftp) (generated by contig assembly with only HiFi data and reference-based scaffolding). Second, TRFill algorithm is used to fill the TOI regions with the help of reference. Third, the assembly of each TOI region is evaluated by the criteria same as the ones used in the experiments on human centromere.

In the experiments of diploid genome, TS2 and TS281 genomes were merged to a simulated diploid genome. The HiFi and Hi-C data of them were also merged to simulate the HiFi and Hi-C data of the diploid genome. Since the merged Hi-C data is lack of the inter-chromosome read-pairs between the original chromosomes from different genomes, sim3C (v0.2)^38^ was used to simulate these Hi-C read pairs which were then added into the Hi-C data. The chromosome-level assembly of the diploid genome is generated by contig assembly with hifiasm (v0.19.5-r587) using only HiFi and Hi-C data and scaffolding with 3dDNA using Hi-C data. The other experimental steps were same with the experiments of haploid genome.

The subtelomeric tandem repeats (TOI regions on query) were identified according to the reference genome. TRF (v4.09.1) was used to detect repetitive patterns within the reference sequences. Then, based on known knowledge, we identify regions located on the edge of each chromosome that harbor monomers with a length of 181bp, and classify them as subtelomeric tandem repeats. Notably, not every chromosome features a pair of subtelomeres due to the absence of 181bp monomers on the edge of chromosomes, and, conversely, some regions containing 181bp monomers are not situated at the chromosome termini. Then the TOI regions on query were identified according to the synteny between reference and query built by Syri (v1.6).

### Population-level Analysis of Tomato Subtelomeric Tandem Repeat Units

The chromosome-level assemblies of 29 tomato genomes were downloaded from the SolOmics database (http://solomics.agis.org.cn/tomato/ftp). The subtelomeric tandem repetitive regions were assembled by TRFill algorithm with the same pipeline as the validation experiments of haploid tomato genome using Heinz1706 genome assembly (assembled in this study using all of HiFi, ONT UL and Hi-C data) as reference. For each subtelomeric tandem repeat region, if the assembly of TRFill algorithm is longer than the original one, the sequence was updated at this region, while the original sequence was kept otherwise.

The population-level analysis was finished with the help of Tandem Repeat Annotation and Structural Hierarchy (TRASH)^39^. TRASH monomer mode and HOR mode were used to detect the monomers and HORs from the tandem repeat sequences respectively, providing the information such as the start and end coordinates, orientation and length of each repeat unit.

In the general analysis of monomers and HORs, a 181bp consensus sequence of monomer was built by multiple sequence alignment of monomers. More specifically, a consensus monomer sequence for each subtelomere was first built according to the multiple sequence alignment of the monomers in this subtelomere generated by an aligner mafft2. Then the global consensus sequence was generated according to the multiple sequence alignment of all subtelomere-level consensus sequences in the same way.

In the sequence-similarity analysis of monomers, comparison was performed 1) between the intra-genome monomers and inter-genome monomers, 2) between intra-chromosome and inter-chromosome monomers in the same genomes, and 3) between intra-subtelomere and inter-subtelomere monomers in the same chromosomes. For each of the comparison, a series of random experiments (a half part for intra and the other half for inter) were done, each of which randomly selected two sets of monomers and the pairwise similarity was calculated between them. The pairwise similarity was defined as the ratio of the number of common monomers (different monomers with completely same sequences) between two sets to the total number of monomers in both sets. For 1) and 2), in each random experiment, each of the two sets contains all monomers in a randomly selected subtelomere. For 3), in each random experiment, each of the two sets contains one tenth of the monomers in a randomly selected subtelomere, and the two sets do not contain same monomers (but may contain different monomers with completely same sequences) for intra-experiments. In the subsequent similarity calculations between all pairs of subtelomeres, the pairwise similarity was defined in the same way.

## Availability

### Code availability

The code of TRFill are available at https://github.com/pants08/TRFill.

### Funding

This work was supported by National Natural Science Foundation of China (Grant No. 32100501); Shenzhen Science and Technology Program (Grant No. RCBS20210609103819020); Innovation Program of Chinese Academy of Agricultural Sciences.

### Author Contributions

H. Wen developed and implemented the TRFill algorithm. W. Pan proposed the idea of reference-guided genome assembly, supervised the project and revised the manuscript.

## Acknowledgements

We thank Haoyu Cheng from Dana-Farber Cancer Institute, Harvard University for discussion about the key idea of this paper. We also thank the funding support for this project.

