## Supplementary Figure for "Reference-guided automatic assembly of genomic tandem repeats with only HiFi and Hi-C data enables population-level analysis"

Weihua Pan:

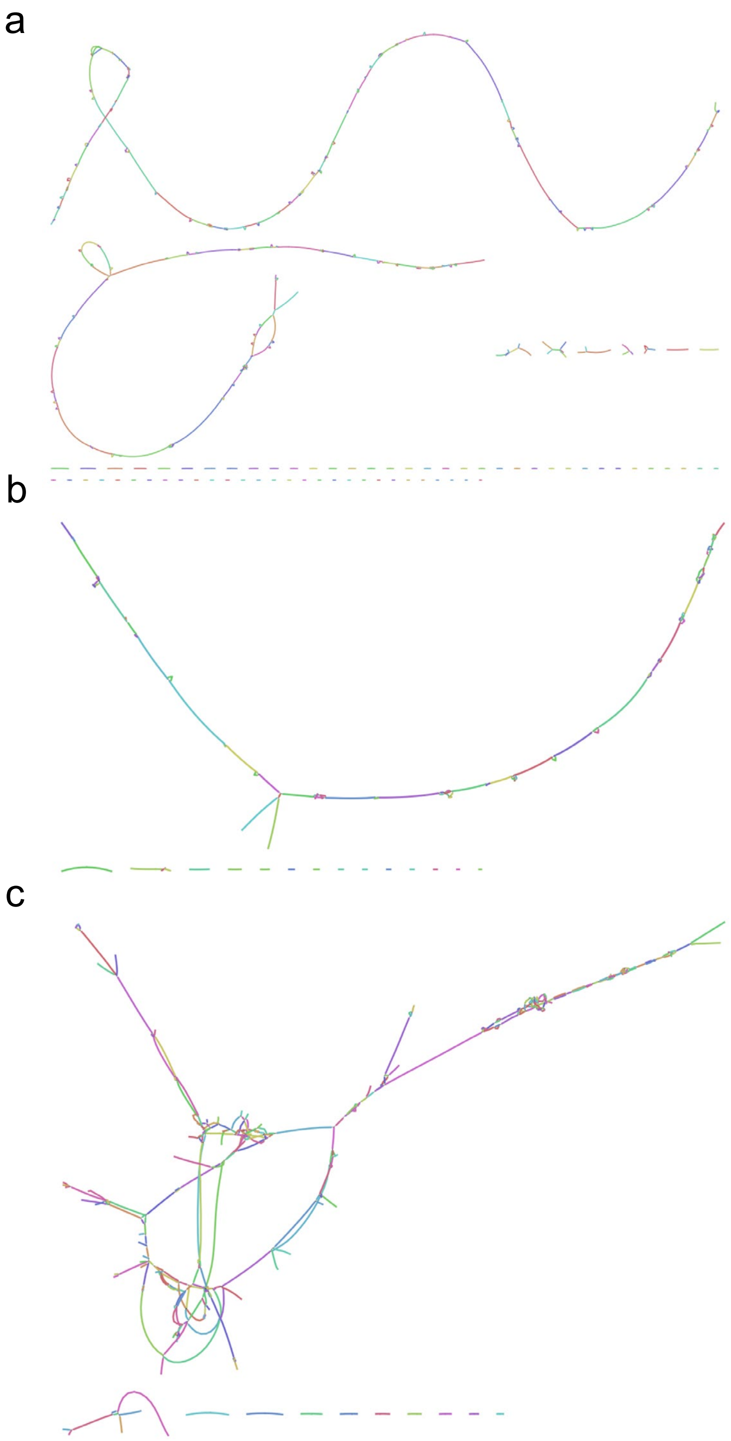

**Supplementary Fig. 1** The unitig graph of HG002 chr1 (**a**), chr16(**b**) and chr6(**c**) output by hifiasm (0.19.5-r587), visualized by Bandage (0.8.1).

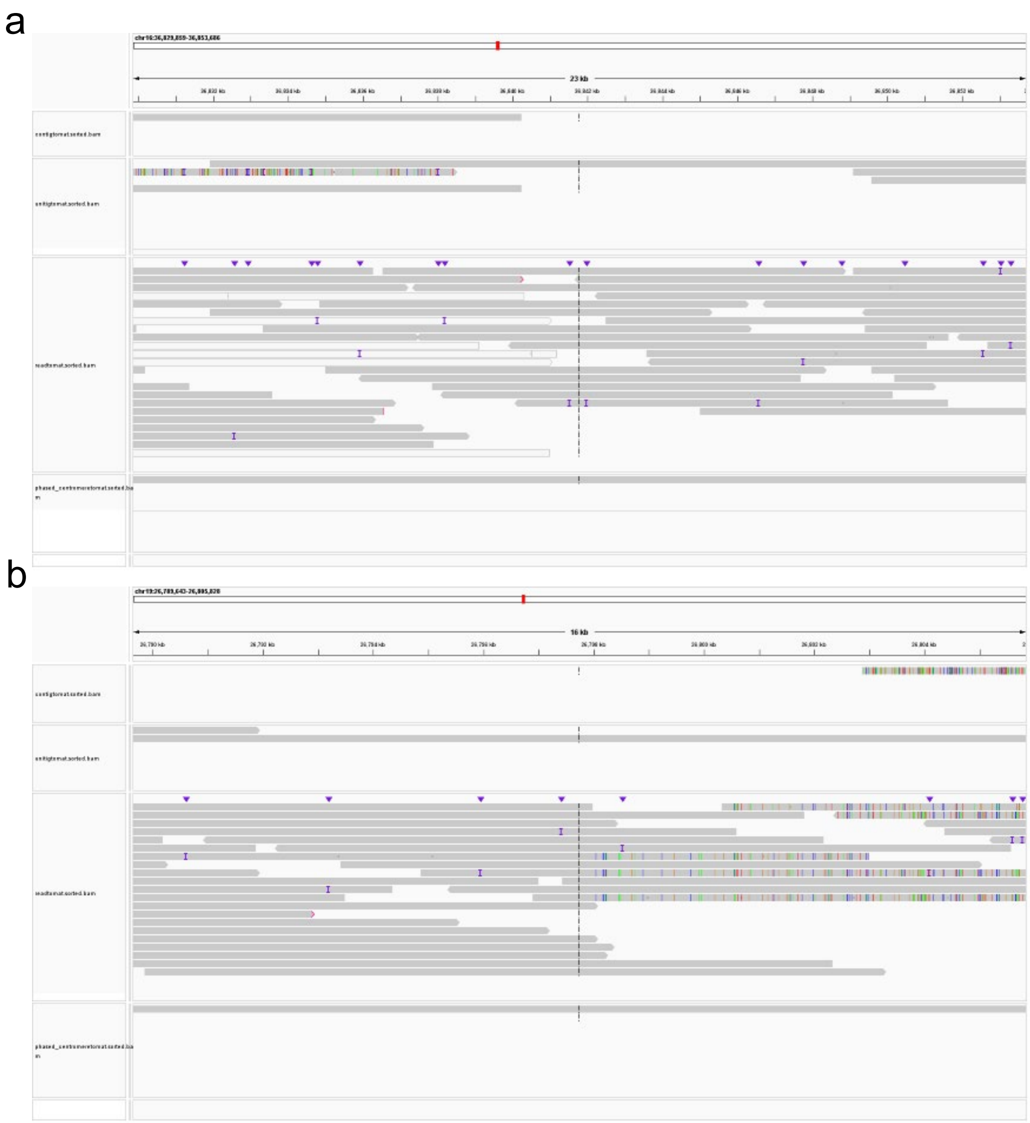

**Supplementary Fig. 2** Visualization of different sequences' alignment to the ground truth assembly with IGV (2.13.2) on HG002 maternal chr16 36,829,859bp~36,853,686bp (**a**) and HG002 maternal chr19 26,789,643~26,805,828bp (**b**). The first row stands for the unitig graph, the second row stands for the contig graph, the third row stands for the reads and the final row stands for corresponding assembly results of TRFill.

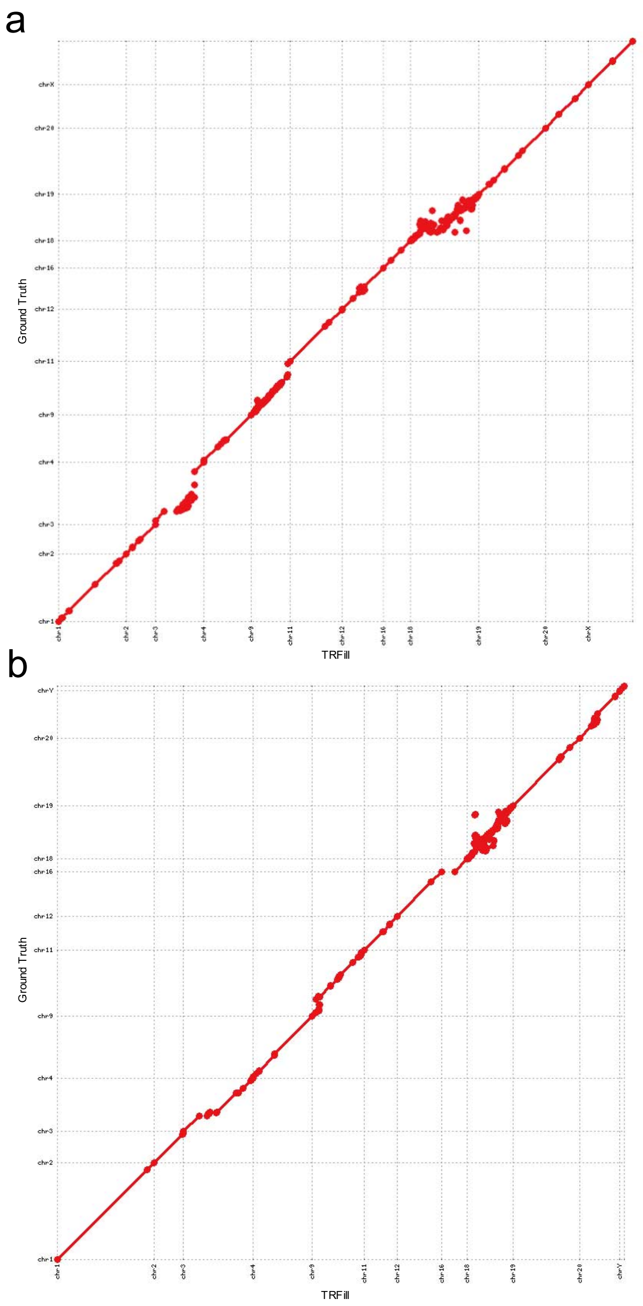


**Supplementary Fig. 3** Synteny plots generated by Mummer between the "ground truth" assemblies and the assemblies after TRFill reassembly of alpha satellite sequences for HG002 maternal (**a**) and paternal (**b**) chromosomes. Only the chromosomes with assemblies obviously improved by TRFill are shown.

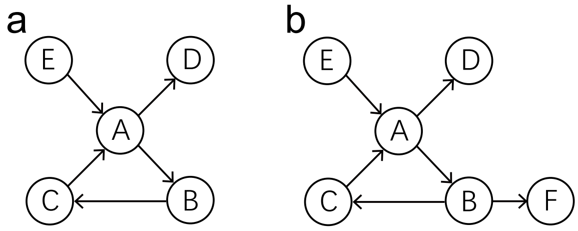


**Supplementary Fig. 4** Illustration of two branching circumstances DFS may encounter during the process of assembling unitigs into contigs. **a**, despite A is a branching node, it is resolvable by revisiting A according to timestamp. Therefore the contig would be generated as $E \rightarrow A \rightarrow B \rightarrow C \rightarrow A \rightarrow D$. **b**, Despite A is resolvable, B is unresolvable since it is a branching node that can not be revisit later. Therefore three contigs would be generated as $E \rightarrow A \rightarrow B, C \rightarrow A \rightarrow D$ and $F$.
